# Task history dictates how the dorsolateral striatum controls action strategy and vigor

**DOI:** 10.1101/2023.01.11.523640

**Authors:** Adam C. G. Crego, Kenneth A. Amaya, Jensen A. Palmer, Kyle S. Smith

## Abstract

The dorsolateral striatum (DLS) is linked to the learning and honing of action routines. However, the DLS is also important for performing behaviors that have been successful in the past. The learning function can be thought of as prospective, helping to plan ongoing actions to be efficient and often optimal. The performance function is more retrospective, helping the animal continue to behave in a way that had worked previously. How the DLS manages this all is curious. What happens when a learned behavior becomes sub-optimal due to environment changes. In this case, the prospective function of the DLS would cause animals to (adaptively) learn and plan more optimal actions. In contrast, the retrospective function would cause animals to (maladaptively) favor the old behavior. Here we find that, during a change in learned task rules, DLS inhibition causes animals to adjust less rapidly to the new task (and to behave less vigorously) in a ‘maladaptive’ way. Yet, when the task is changed back to the initially learned rules, DLS inhibition instead causes a rapid and vigorous adjustment of behavior in an ‘adaptive’ way. These results show that inhibiting the DLS biases behavior towards initially acquired strategies, implying a more retrospective outlook in action selection when the DLS is offline. Thus, an active DLS could encourage planning and learning action routines more prospectively. Moreover, the DLS control over behavior can appear to be either advantageous/flexible or disadvantageous/inflexible depending on task context, and its control over vigor can change depending on task context.

**Significant Statement:** Basal ganglia networks aid behavioral learning (a prospective planning function) but also favor the use of old behaviors (a retrospective performance function), making it unclear what happens when learned behaviors become suboptimal. Here we inhibit the dorsolateral striatum (DLS) as animals encounter a change in task rules, and again when they shift back to those learned task rules. DLS inhibition reduces adjustment to new task rules (and reduces behavioral vigor), but it increases adjustment back to the initially learned task rules later (and increases vigor). Thus, in both cases, DLS inhibition favored the use of the initially learned behavioral strategy, which could appear either maladaptive or adaptive. We suggest that the DLS might promote a prospective orientation of action control.

## Introduction

The dorsolateral striatum (DLS) is necessary for invigorating actions, organizing them as sequences, and expressing them as habits (Amaya & Smith, 2018; Barker & Taylor, 2014; Corbit & Janak, 2016; Gasbarri et al., 2014; Graybiel, 2008; Knowlton & Patterson, 2018; Lerner, 2020; Lovinger & Gremel, 2021; Malvaez & Wassum, 2018; Seger, 2018; Yin & Knowlton, 2004). Accentuation of DLS activity during the initiation of a behavior is linked with how vigorous that behavior is (Barnes, et al., 2005; Crego et al., 2020; Jin & Costa, 2010; Jog et al., 1999; Regier et al., 2015; Smith & Graybiel, 2013a; Yttri & Dudman, 2016). The DLS is particularly important for causing extensively trained behaviors to be continued, by habit, when the outcome is delivered noncontingently from the action or when the outcome is devalued (Yin et al., 2004, 2006). DLS activity would thus seem to promote the performance of action routines that have worked previously. In line with this, neural representations of reward locations within the DLS are found to favor previously experienced locations, rather than forthcoming locations (Cunningham et al., 2021). This function of the DLS can be regarded as carrying an outlook that is retrospective and focused on continuing learned behavioral patterns.

However, the DLS is also implicated in action learning and in the honing of ongoing behavior (e.g., (Bailey & Mair, 2006; Bergstrom et al., 2018; Goodman et al., 2017; Packard & McGaugh, 1992; Turner et al.,, 2022; Yin et al., 2009; Yu et al,, 2009)), suggesting a possible role for the DLS in the prospective planning of actions. Also, because many studies examine DLS activity when task conditions are stable, it becomes impossible to know if manipulations of the DLS that reduce an aspect of performance do so because planning is affected or prior learning is affected. Moreover, without details on how animals are behaving when behaviors persist in the face of changed action-reward relationships, it can be unclear if they are actually stuck in previously learned routines or if they are exploring the new task conditions and learning (Bailey & Mair, 2006; Dezfouli & Balleine, 2012; Tolman, 1932).

One way to examine how the DLS regulates prospective vs. retrospective action control is to pit these two possibilities against one another. If DLS activity promotes exploiting an already-learned behavior, then DLS inhibition should cause animals to adapt to a change in task conditions more readily. If instead DLS activity promotes exploring and adapting of actions to the current environment, then DLS inhibition should do the opposite and cause animals to be stuck in what they learned and slow to adopt the new task rules. In a recent study (Amaya & Smith, 2021), we did this and found that inhibition of DLS cholinergic interneurons led to more rapid adaptation to changed task rules, which suggested that these neurons promote the performance of learned behaviors (i.e., a retrospective action function). How this relates to the rest of the DLS neuronal population is unclear.

Separately, the DLS helps animals perform actions in a fluid and vigorous manner. Disruption of the DLS or its dopaminergic input causes slower and more variable performance patterns (Bailey & Mair, 2006; Crego et al., 2020; Cromwell & Berridge, 1996; da Silva et al., 2018; Graybiel, 2008; Panigrahi et al., 2015; Yin et al., 2004; Yttri & Dudman, 2016). Stimulating the DLS, particularly at the onset of actions, can directly increase performance vigor as well (Crego et al., 2020). Whether DLS contributes to action vigor in a manner that serves behavioral exploitation (retrospective) or exploration (prospective) is unclear. Thus, when DLS activity is engaged, it could be that animals are more vigorously performing what they have previously done or that animals are attuned to the current environment and are vigorously performing actions that best suit it. Here, we attempt to better understand prospective vs. retrospective DLS functions and the role of performance vigor within those functions.

## Materials and Methods

We incorporated an optogenetic strategy to inhibit the DLS after animals learned a task and encountered surprising changes in the action-outcome relationship. We allowed animals to perform in a self-directed manner in a free operant environment, and we used a battery of behavioral measures to understand what aspects of their behavior and their performance strategies did and did not change during DLS inhibition when the task rules shifted. One important aspect of this design is that animals could continue to use their previously learned behavioral strategy to get reward, but with the shift in task rules that strategy was no longer the most optimal for efficient performance. Shifts in task rules away from a learned strategy, and then back towards that strategy, were employed.

### Subjects and Surgery

Male Long Evans rats (n = 18) were housed individually in a dedicated animal vivarium. All procedures were approved by the Dartmouth College Institutional Animal Care and Use Committee. All rats used were at 250-400 g and were maintained on an 85% post-surgical weight for training and testing. Rats were housed on a reverse light-dark cycle, with all operant chamber experiments conducted in the dark with a house light on in sound attenuated boxes. Surgeries were performed using aseptic techniques under isoflurane anesthesia: intracranial injection of viral vectors, followed by intracranial implantation of fiber optic guides. Rats received bilateral injections (0.3 µL) into DLS of a single viral construct: eNpHR3.0/halorhodopsin (n = 9, AAV5-hSYN-eNpHR3.0-EYFP) or control (n = 9, AAV5-hSYN-EYFP) using a 33-gauge syringe and microinfusion pump. Bilateral DLS coordinates for viral injections were, in mm: AP +0.5, ML −/+ 4.0, DV injection −4.3 from skull, with fiber implants (200 µm, ThorLabs or in-house) terminating at DV −3.8 mm. Fiber implants were permanently affixed to the skull using dental cement and skull screws. All rats received at least one-week post-surgery time to recover.

### Experimental Setup

Rats underwent training in eight identical modular operant chambers (Med Associates) in a dedicated experimental room. All chambers were housed in their own sound attenuated box and closed during all training sessions. Chambers were equipped with two identical levers that were retractable. Both levers were inserted upon the start of a session and remained inserted until the completion of the session. For each animal, one lever was active (paired with reward) and the other inactive. In between the levers was a magazine food cup receptacle, where sucrose reward pellets would drop upon the specified Med PC macro script program. At the top back region of the chambers, a house light would illuminate to indicate a session had begun and remain on for the duration of the experiment, but the remainder of the testing room was in the dark. A photobeam within the magazine acted as a reset when quantifying specific measures of lever pressing (e.g., inter-trial-intervals, lever press bouts, etc.). All sessions began with house light illumination and the insertion of two levers. All training and experimental sessions would end after 60 minutes or once 30 rewards were earned, with the exception of a RT30 (random time) day always ending in 30 minutes.

### Optogenetic Manipulations

Optical patch cords with a 0.2 µm fiber core and 0.39 NA were used for light administration (Thorlabs, Newton, NJ; Doric Lenses, Quebec City, Canada). On days where DLS was manipulated, a 593.5 nm light from DPSS lasers (Shanghai Laser & Optics Century Co., Ltd.) was delivered through these fiber patch cords that were connected to a rotary-joint beam splitter (Thorlabs, Newton, NJ; Doric Lenses, Quebec City, Canada). This allowed for two fiber patch cables to be connected directly to the surgically implanted optic fiber cannula (in-house fiber implants, 200 µm, ThorLabs) by ceramic sleeves. Laser illumination delivery was gated and triggered by TTL pulses through designed Med PC programs, to individual operant chamber boxes. Power output was measured as 3-5 mW directly from each patch cable ferrule connector prior to and after laser test days. Fiber implants were also tested post-perfusion to confirm their patency.

Rats were habituated to the optogenetic cables prior to any experimental days. On separate manipulation days, laser light was delivered into the DLS as a continuous 1-sec pulse triggered by the inflection of a pre-defined paired reward lever press. The 1-sec illumination duration was chosen to allow animals to behave as they would during that 1-sec period, and 1-sec captured well a typical duration between lever deflection and magazine entry. Any unpaired lever pressing did not result in laser illumination delivery. Operant behavior and task events (e.g., lever presses, time, etc.) were recorded by Med PC software via a designated computer. Accuracy of automated behavioral measurements was verified through videotaping and hand scoring of a subset of sessions.

### Operant Chamber Task

Rats were first trained on a fixed-ratio 1 (FR1) schedule of reward, where a predefined active lever (i.e., consistent throughout the duration of the experiment) was paired with reward delivery. An inactive lever was paired with nothing. Lever assignment was counterbalanced across groups. FR1 training sessions were administered for 7 sequential days, wherein rats learned a contingency that a single lever press resulted in the delivery of a single reward pellet within the magazine port. This learned behavior was then challenged by a change in the action requirement for the reward, which was a surprise upshift in requirement from one to three presses for reward (FR1 → FR3; “upshift day”). This sudden shift now required rats to press the previously assigned paired-rewarded lever three times. The DLS was optogenetically inhibited (eNpHR3.0) on only the first press of the active lever during this upshift day.

A series of 7 FR3 training days followed without further DLS inhibition to evaluate how animals adjusted to the new task rules. Then, the DLS was inhibited during a FR3 “maintenance day”. This day was conducted identically to the upshift day, and it was followed by a single non-illumination FR3 day for comparison. This maintenance day was meant to provide a comparison of how DLS inhibition affected performance when the FR3 task rule was new versus established/unchanged.

Following this, the task was then suddenly returned back to the originally learned FR1 schedule (FR3 → FR1; “downshift day”). DLS illumination was time-locked to the first press after a magazine entry (i.e., subsequent pressing would not yield illumination until a magazine entry occurred again). This downshift day allowed us to understand the effects of DLS inhibition when FR1-based performance was the learned rule to adapt against (upshift day) versus when it was a new rule to adapt towards (downshift day).

A series of 7 FR1 training days followed without DLS inhibition to again establish performance on that FR1 schedule. Then, and finally, rats were subjected to a surprise introduction of a random-time 30 (RT30) schedule in which reward was delivered entirely independently from any pressing. This shift to RT30 was designed to entirely remove the action requirement for reward, allowing us to examine how DLS inhibition might affect the frequency, distribution, and pattern of learned actions when they were no longer necessary. This procedure is similar to one we recently published to probe behavioral exploration vs. behavioral fixity in understanding DLS function (Amaya & Smith, 2021) (see also: (Balleine & Killcross, 1994; Balleine, Killcross, & Dickinson, 2003)). Illumination was applied during this RT30 period, and it was paired with every lever press (i.e., previously paired-rewarded lever) regardless of magazine entries.

### Histology

At the termination of each cohort, lethal doses of anesthesia (sodium pentobarbital) were administered. This was followed by a transcardial perfusion of 0.9% saline and 4% paraformaldehyde. Brains were fixed further in 20% sucrose solution overnight and frozen at −80 °C. Brains were sectioned at 30-60 µm using a microtome, and later mounted to slides and cover slipped with a DAPI-containing medium. Fiber placement and EYFP expressing neuron spread were analyzed using a fluorescent microscope (Olympus BX53 fluorescent microscope with DP73 camera).

### Data Analysis

Lever deflections and pellet magazine entries were recorded through Med PC. Operant behavioral measures were individually extracted from Med PC files using a custom Python script. Statistical testing was conducted using R (R Core Team, 2016). All graphs were made using R (R; “ggplot2”) and stylized in Adobe Illustrator.

Individual generalized linear mixed models (R; “lme4”) were created for each dependent behavioral variable (Gamma distribution for temporal measures: overall time (min), inter-press intervals (sec), reward contact to lever press (sec), lever press to mag entry (sec); Binomial distribution for action measures averaged: 1-press bouts, and 3-press bouts) for within- and between-subject analyses for main effects day (illumination day and subsequent non-laser day), virus group (control vs. eNpHR3.0), and the interaction between these variables. These comparisons were conducted separately based on *a priori* events of interest for DLS function on separate illumination days (e.g., upshift, maintenance, downshift, RT30). Reported statistics include parameter estimates, 95% confidence intervals, and P-values (R; “lmerTest”, (Kuznetsova, Brockhoff, & Christensen, 2017)). Separately, a linear mixed model was created to analyze the effects of day, group, and their interaction on presses made per reward. Linear mixed models were fit by maximum likelihood and t-tests used Satterthwaite approximations of degrees of freedom (R; “lmerMod”). Linear models were analyzed with package lme4 from CRAN (Bates et al., 2015). Reported statistics include parameter estimates (β values), 95% confidence intervals, and P-values.

For behavior during the RT30 phase of the experiment, measures of action probability just prior to and just following reward delivery were analyzed. Time windows for these analyses were individually set as the mean inter-press interval, rounded up. For example, if an animal averaged 2.3s between presses, the pre- and post-reward delivery windows were set as 3s before and 3s after reward delivery. Models of action probability were generalized linear mixed models with fixed effects of group, session, and their interaction and random intercepts for individual start points on session 1. Reported statistics include odds ratios, 95% confidence intervals, and P-values. Lever press rates (as presses per minute) as predicted by group assignment and session were assessed both overall and within a peri-reward time window (+/-1 second). Linear mixed models with fixed effects of group, session, and their interaction and random intercepts for individual rats were used here to predict respective dependent variables, either overall presses per minute or presses per minute within the specified time window. Lever press rates and action probabilities were then assessed with respect to laser delivery, with lever press rates outside of laser delivery being considered distinct from lever press rates during laser delivery. Action probability was calculated as the probability of a lever press occurring during laser administration as a simple binary (yes/no) as each lever press triggered laser delivery during this phase of the experiment. A linear mixed model with fixed effects of laser bin, group, session, and interaction terms with random intercepts for individual rats was used to predict presses per minute made. Importantly, Session was re-centered such that the final RT-30 session is the comparison point for the main effects of group and laser bin. To model action probability during laser delivery, a generalized linear mixed model with fixed effects of group, session, and their interaction with random intercepts for individual rat starting points was used to predict probability of action. Reported statistics include parameter estimates (or odds ratios for GLMMs), 95% confidence intervals, and P-values.

## Results

Rats were trained and tested in a free operant conditioning environment (Fig. 1A) to investigate DLS function of performing a learned lever-press action for a reward outcome (Balleine et al., 2009). The DLS was inhibited online via light-gated halorhodopsin on a series of test days. During these test days new task rules were imposed: upshift (FR1→FR3 rule change), maintenance FR3 day (for comparison), downshift (FR3→FR1), and random time (FR1→RT30) (Fig. 1A). Effects of DLS inhibition were compared to a control group receiving identical treatment but lacking halorhodopsin expression. Histology confirmed DLS opsin expression (Fig. 1B) and fiber placements (Fig. 1C) in both groups (Fig. 1D). Previous work (Crego et al., 2020) showed that roughly 83% of neurons in the DLS express these viral constructs.

**Figure 1.**
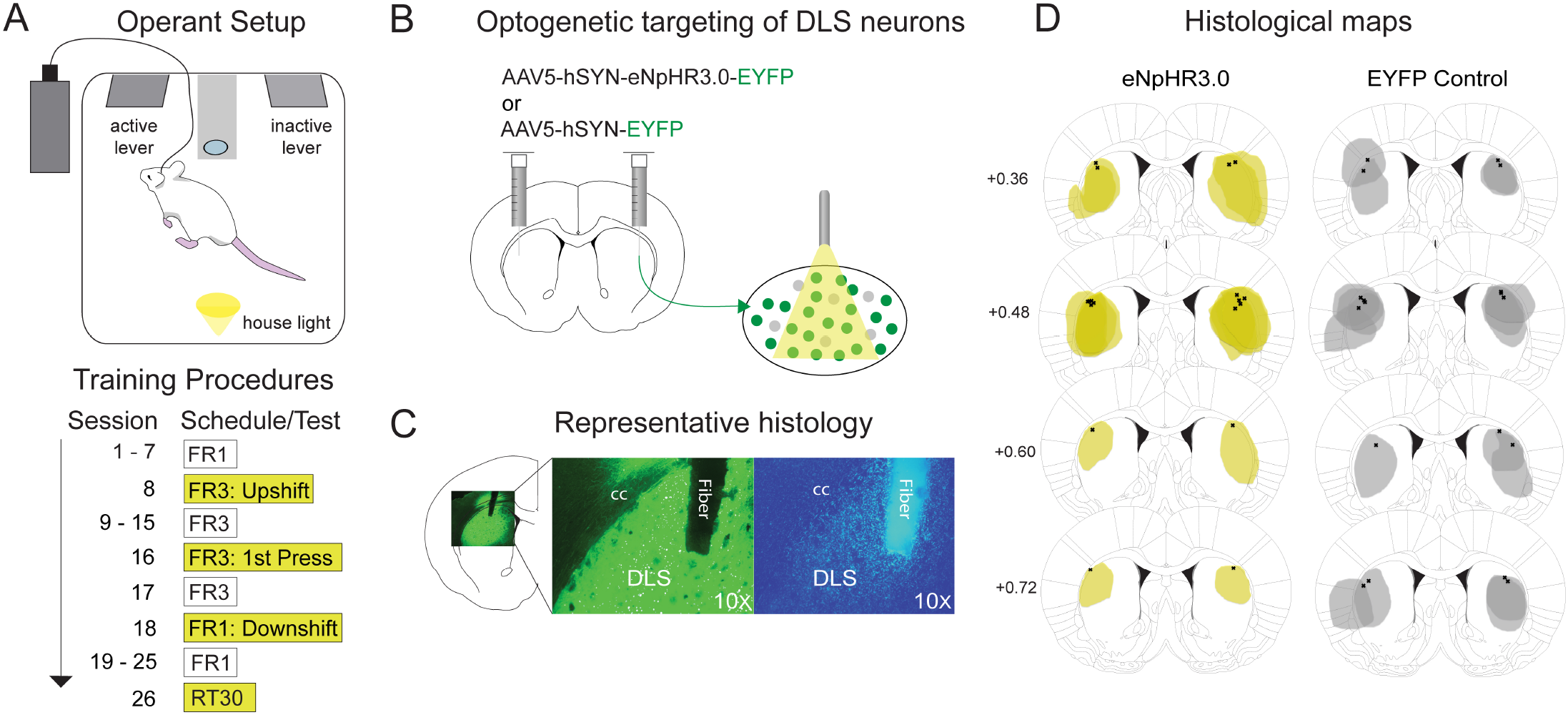
Operant chamber task. **A**, Top, operant task cartoon illustrating arrangement of the magazine reward port nestled between the active lever (i.e., rewarded) and inactive lever (i.e., unrewarded). Optogenetic setup was external to the chamber, with cables threaded through the top. House light illumination signaled the start of a session. Bottom, training protocol and four illumination test days: contingency tests: Upshift (FR1 → FR3), Downshift (FR3 → FR1), and three RT30 sessions; maintenance test: illumination on 1^st^ press after FR3 training. **B**, Cartoon representation of bilateral injections of viral constructs into DLS; green dots = EYFP expression, grey dots = no expression. **C**, Example optogenetic expression of DLS section from left to right: EYFP and DAPI fluorescent stain for neuronal health. Fiber placement labeled across sections. **D**, Expression areas (left to right) of eNpHR3.0-EYFP and EYFP control across animals. Expression is denoted per animals as semi-transparent shading overlaid across rats and group. Numbers on the right denote anteroposterior plane in mm relative to bregma. Outside DLS expression was observed; however, fiber placements denoted as X’s suggested illumination targets were exclusive to DLS. FR, fixed-ratio, RT, random-time. EYFP, enhanced yellow fluorescent protein. DAPI, 4’,6’-diamidino-2-phenylindole. eNpHR3.0, halorhodopsin. cc, corpus collosum; DLS, dorsolateral striatum.

### Upshift Day (FR1→ FR3)

#### Temporal data

On the upshift day rats were switched from FR1 to FR3 with DLS inhibition occurring on the first press of the action sequence. The overall time to complete the session was slower for both groups compared to the subsequent non-illumination FR3 day, and within the illumination upshift day the halorhodopsin group was slower than the control group (Fig. 2A). This was shown by a significant interaction between day and group (estimate: 0.79, CI: 0.68 – 0.92, *p* = 0.002), a main effect of day (estimate: 0.63, CI: 0.54 – 0.73, *p* < 0.001), and no effect of subject group (estimate: 1.27, CI: 0.95 – 1.70, *p* = 0.107). As contributing factors to overall task completion time, the time rats spent between lever presses (i.e., inter-press intervals) were greater for halorhodopsin rats on the upshift day (Fig. 2B) as shown by a significant interaction between day and group (estimate: 0.89, CI: 0.82 – 0.96, *p* = 0.003), and a main effect of day (estimate: 0.78, CI: 0.72 – 0.85, *p* < 0.001) but not group (estimate: 1.14, CI: 0.86 – 1.53, *p* = 0.363). Similarly, latencies between a magazine entry and the next lever press (Fig. 2C) showed a main effect of day (estimate: 0.74, CI: 0.66 – 0.82, *p* < 0.001) and a significant interaction between day and group (estimate: 0.80, CI: 0.72 – 0.88, *p* < 0.001), but no effect of group (estimate: 0.89, CI: 0.63 – 1.27, *p* = 0.524), indicating slower magazine-to-press rates during DLS inhibition. Lastly, we analyzed the latency between the last FR3 press and the subsequent reward contact (Fig. 2D) within the magazine, where there was a main effect of day (estimate: 0.90, CI: 0.86 – 0.94, *p* < 0.001) but no effect of group (estimate: 1.09, CI: 0.89 – 1.34, *p* = 0.406), and there was not an interaction between day and group (estimate: 0.98, CI: 0.93 – 1.02, *p* = 0.319). Thus in general, DLS inhibition led to a slowing of behavior chiefly between presses and between a magazine entry and a press but reward retrieval was unimpacted.

**Figure 2.**
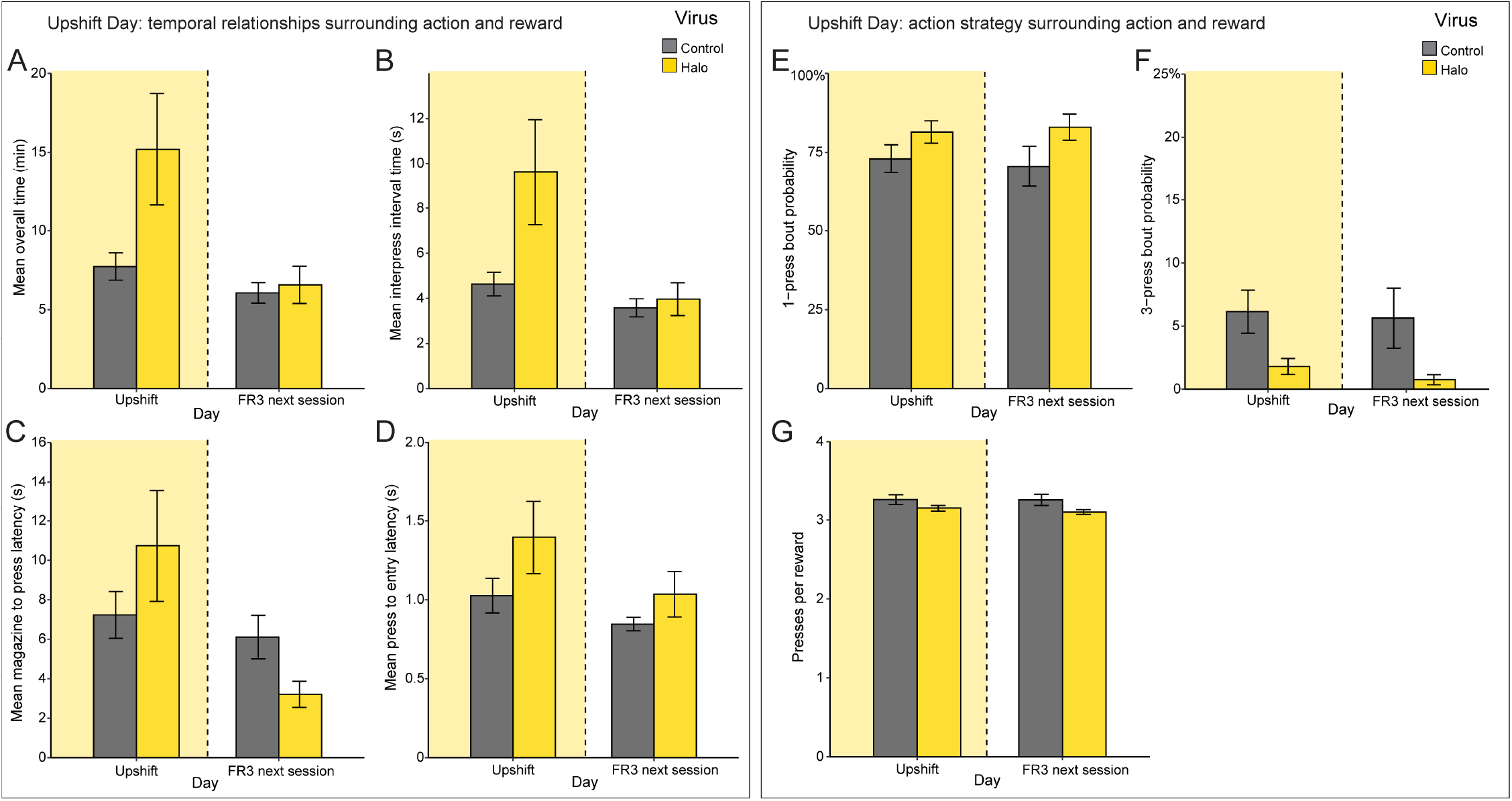
Upshift illumination day. 1-sec continuous DLS illumination on the first press of a FR3 sequence. Illumination effects were compared within- and between-groups to a normal FR3 session the day immediately after. **A**, Mean overall time (min). **B**, Mean inter-press intervals (sec). **C**, Mean reward contact (magazine) to press latency (sec). **D**, Mean lever press to a magazine entry (sec). **E**, Mean 1-press bout probability (%). **F**, Mean 3-press bout probability (%). **G**, Total presses per reward (ratio). Control, grey; eNpHR3.0, yellow. For all figure panels, bars and errors show mean ± SEM; asterisks denote significant comparisons (\**P*<0.05, \*\**P*<0.01, \*\*\**P*<0.001); lack of asterisks denotes lack of significance.

#### Action strategy data

Animals can in principle display a range of action patterns reflective of an underlying strategy, which we assessed by examining pressing bout probability, presses per reward, and rate of pressing surrounding reward. On the upshift day, as well as on the subsequent day, the majority of actions occurred in 1-press bouts (i.e., press then check the magazine before pressing again), which had been optimal under the learned FR1 conditions (Fig. 2E). For 1-press bouts, there was no day effect (estimate: 0.95, CI: 0.83 – 1.08, *p* = 0.428), no day and group interaction (estimate: 1.12, CI: 0.92 – 1.36, *p* = 0.246), and no group effect (estimate: 1.89, CI: 0.89 – 4.05, *p* = 0.099). Thus, while the halorhodopsin group trended towards performing more 1-press bouts, this was not significant. The theoretically optimal strategy during FR3 task conditions is to perform 3-press bouts (i.e., press 3 times before checking the magazine). On the illumination and subsequent days, these 3-press bouts were low in number and there was no effect of day (estimate: 0.93, CI: 0.72 – 1.19, *p* = 0.546) or day and group interaction (estimate: 0.69, CI: 0.38 – 1.23, *p* = 0.203) (Fig. 2F). However, there was a main between groups effect (estimate: 0.19, CI: 0.06 – 0.61, *p* = 0.005) due to the halorhodopsin group performing significantly fewer 3-press bouts on the illumination day, which appeared to carry over to the subsequent day as well.

Further analysis on the number of lever presses performed per reward (Fig. 2G) revealed no main effect of day (estimate: −0.01, CI: −0.05 – −0.02, *p* = 0.392) or day and group interaction (estimate: −0.01, CI: − 0.04 – −0.02, *p* = 0.458). Similar to the 3-bout results, however, there was a main between groups effect (estimate: −0.07, CI: −0.13 – −0.01, *p* = 0.027) reflecting fewer presses per reward in the halorhodopsin group. To summarize, DLS inhibition led to a less vigorous set of behavior and tended to bias animals towards a more task-optimal FR3 action strategy.

### FR3 Maintenance Day

#### Temporal data

After the upshift illumination day, all rats were trained under the new FR3 requirement for the next seven days. To inform a possible mechanism for DLS in ongoing action performance (i.e., when there is no change in the action requirement for reward), a maintenance FR3 illumination day was conducted during which DLS was inhibited. For task completion time, there was a main effect of day (estimate: 0.79, CI: 0.68 – 0.92, *p* = 0.003) and a day/group interaction (estimate: 0.73, CI: 0.63 – 0.86, *p* < 0.001), but not a group effect (estimate: 1.11, CI: 0.81 – 1.52, *p* = 0.534) (Fig. 3A). Halorhodopsin animals were thus slightly slower on the illumination day, while if anything slightly faster on the subsequent day. Inter-press intervals (Fig. 3B) followed the same trend, with main effects of day (estimate: 0.90, CI: 0.83 – 0.96, *p* = 0.003) and a day/group interaction (estimate: 0.92, CI: 0.85 – 0.98, *p* = 0.016), but no group effect (estimate: 0.93, CI: 0.68 – 1.27, *p* = 0.639). Likewise, the press to magazine entry latency (Fig. 3D) had a day and group interaction (estimate: 0.91, CI: 0.85 – 0.98, *p* = 0.018) but not an effect of group (estimate: 1.01, CI: 0.85 – 1.21, *p* = 0.877) or day (estimate: 0.92, CI: 0.85 – 0.99, *p* = 0.031). Dissimilar to the upshift illumination day was experimental rats’ quicker ability to go from the magazine to the next press (Fig. 3C) as shown by a main day and group interaction (estimate: 0.93, CI: 0.88 – 0.98, *p* = 0.012); however, neither day (estimate: 0.96, CI: 0.91 – 1.02, *p* = 0.157) or group (estimate: 0.73, CI: 0.49 – 1.08, *p* = 0.117) showed the same effect. In short, there were some rather inconsistent changes in task performance times due to DLS inhibition, with the overall direction in these changes similar to, but not identical to, the changes observed during the upshift test day.

**Figure 3.**
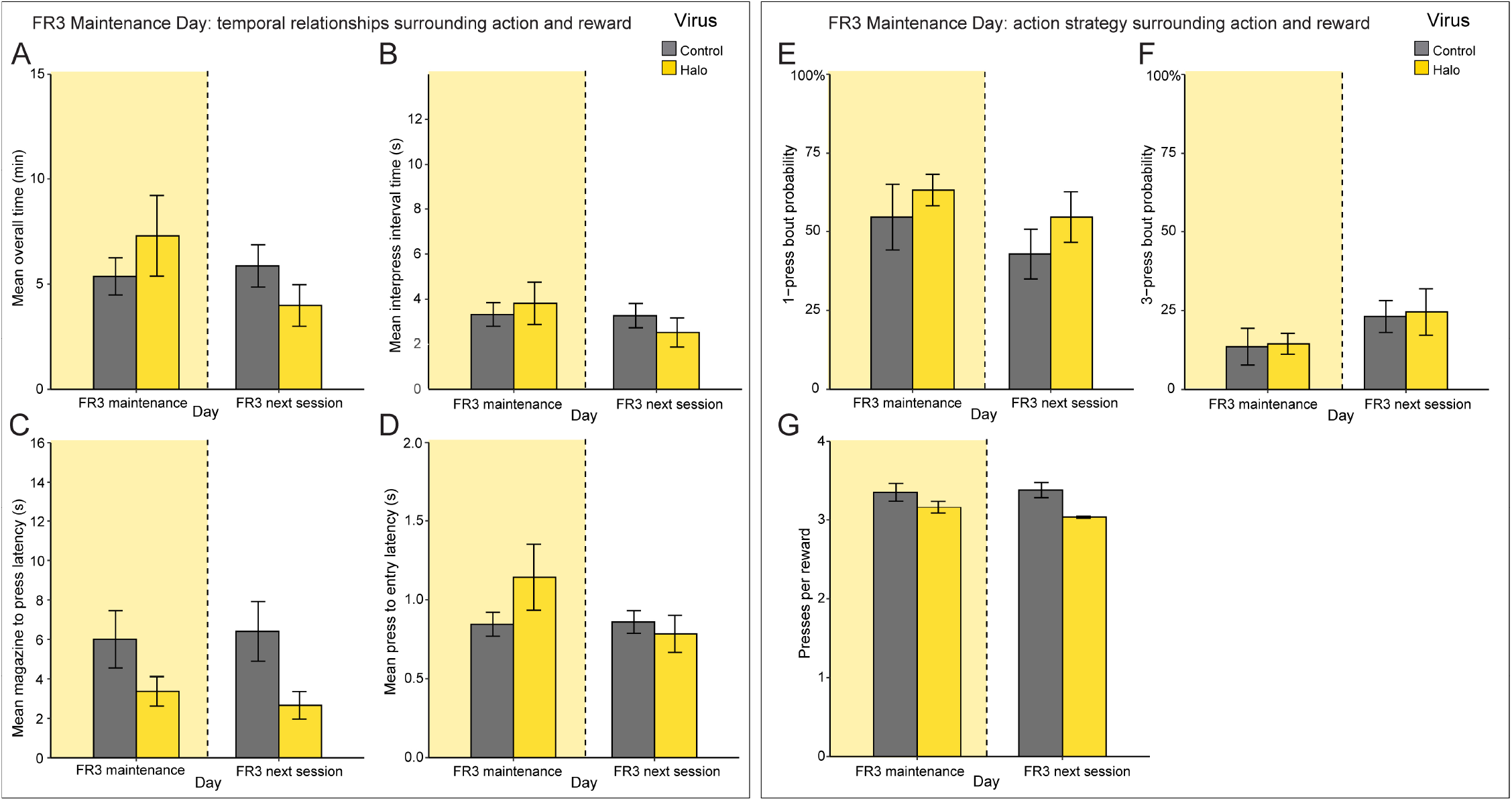
FR3 maintenance illumination. 1-sec continuous DLS illumination on the first press of a FR3 sequence. Illumination effects were compared within- and between-groups to a normal FR3 session the day immediately after. **A**, Mean overall time (min). **B**, Mean inter-press intervals (sec). **C**, Mean reward contact (magazine) to press latency (sec). **D**, Mean lever press to a magazine entry (sec). **E**, Mean 1-press bout probability (%). **F**, Mean 3-press bout probability (%). **G**, Total presses per reward (ratio). Control, grey; eNpHR3.0, yellow. For all figure panels, bars and errors show mean ± SEM; asterisks denote significant comparisons (\**P*<0.05, \*\**P*<0.01, \*\*\**P*<0.001); lack of asterisks denotes lack of significance.

#### Action strategy data

After the 7 FR3 training days, animals still exhibited a mostly FR1-like strategy of behavior, but nevertheless had moved further in the direction of behaving in alignment with the FR3 rule. Thus, 1-press bouts were still dominant but tended to be less numerous than the initial FR3 experience (upshift day) while 3-press bouts showed the reverse trend. On this maintenance day, the increase in 1-press bouts in halorhodopsin rats that was a trend during the upshift day was significant (Fig. 3E) as indicated by a day and group interaction (estimate: 1.51, CI: 1.11 – 2.07, *p* = 0.009) and group (estimate: 2.76, CI: 1.16 – 6.56, *p* = 0.022), but not day (estimate: 1.04, CI: 0.86 – 1.26, *p* = 0.690). 3-press bouts seemed less affected on this day by DLS perturbation (Fig. 3F): there was no main effect of group (estimate: 0.37, CI: 0.12 – 1.13, *p* = 0.082), day (estimate: 1.00, CI: 0.72 – 1.39, *p* = 1.000), or a day and group interaction (estimate: 0.56, CI: 0.29 – 1.09, *p* = 0.091).

Presses per reward (Fig. 3G) showed similar results as in the upshift day with a main effect of day (estimate: −0.09, CI: −0.15 – −0.02, *p* = 0.015) and group (estimate: −0.35, CI: −0.64 – −0.05, *p* = 0.021), but not a day and group interaction (estimate: 0.01, CI: −0.09 – 0.11, *p* = 0.822). In summary, DLS inhibition created modest and somewhat inconstant changes in performance vigor, and likewise led to a modest and somewhat inconsistent trend towards more FR1-like behavior.

### Downshift Day (FR3→ FR1)

#### Temporal data

In the next stage of the experiment, the task rules were again suddenly changed from FR3 back to FR1 and the DLS was inhibited. Thus, rats could ‘re-use’ the initially learned and familiar 1-press bout strategy. Strikingly, rats with DLS inhibition were exceptionally fast in their completion of the task on this day (Fig. 4A). There were significant effects of day (estimate: 0.88, CI: 0.83 – 0.92, *p* < 0.001) and a day/group interaction (estimate: 0.90, CI: 0.85 – 0.95, *p* < 0.001). There was no group effect (estimate: 0.87, CI: 0.66 – 1.16, *p* = 0.349). Mean inter-press intervals followed a similar trend (Fig. 4B). While there were no effects of day (estimate: 1.02, CI: 0.96 – 1.08, *p* = 0.467) or group (estimate: 0.68, CI: 0.34 – 1.37, *p* = 0.281), there was a significant day/group interaction (estimate: 0.91, CI: 0.83 – 0.98, *p* = 0.018). Latency between magazine entries and the next press (Fig. 4C) revealed no main effect of day (estimate: 1.00, CI: 0.95 – 1.06, *p* = 0.882); however, we did find significant effects for group (estimate: 0.44, CI: 0.22 – 0.89, *p* = 0.023) and a day/group interaction (estimate: 0.86, CI: 0.80 – 0.93, *p* < 0.001). Lastly, mean time between a press and subsequent reward contact (Fig. 4D), did not show any effects of day (estimate: 0.93, CI: 0.84 – 1.02, *p* = 0.126), group (estimate: 1.11, CI: 0.81 – 1.51, *p* = 0.525), or a day/group interaction (estimate: 0.99, CI: 0.86 – 1.13, *p* = 0.865). Thus, the robustly sped-up task completion time in halorhodopsin rats reflected quicker time between presses and quicker time to transition from magazine entry back to pressing. These were the same task moments that were most affected (and oppositely affected) by DLS inhibition during the upshift test.

**Figure 4.**
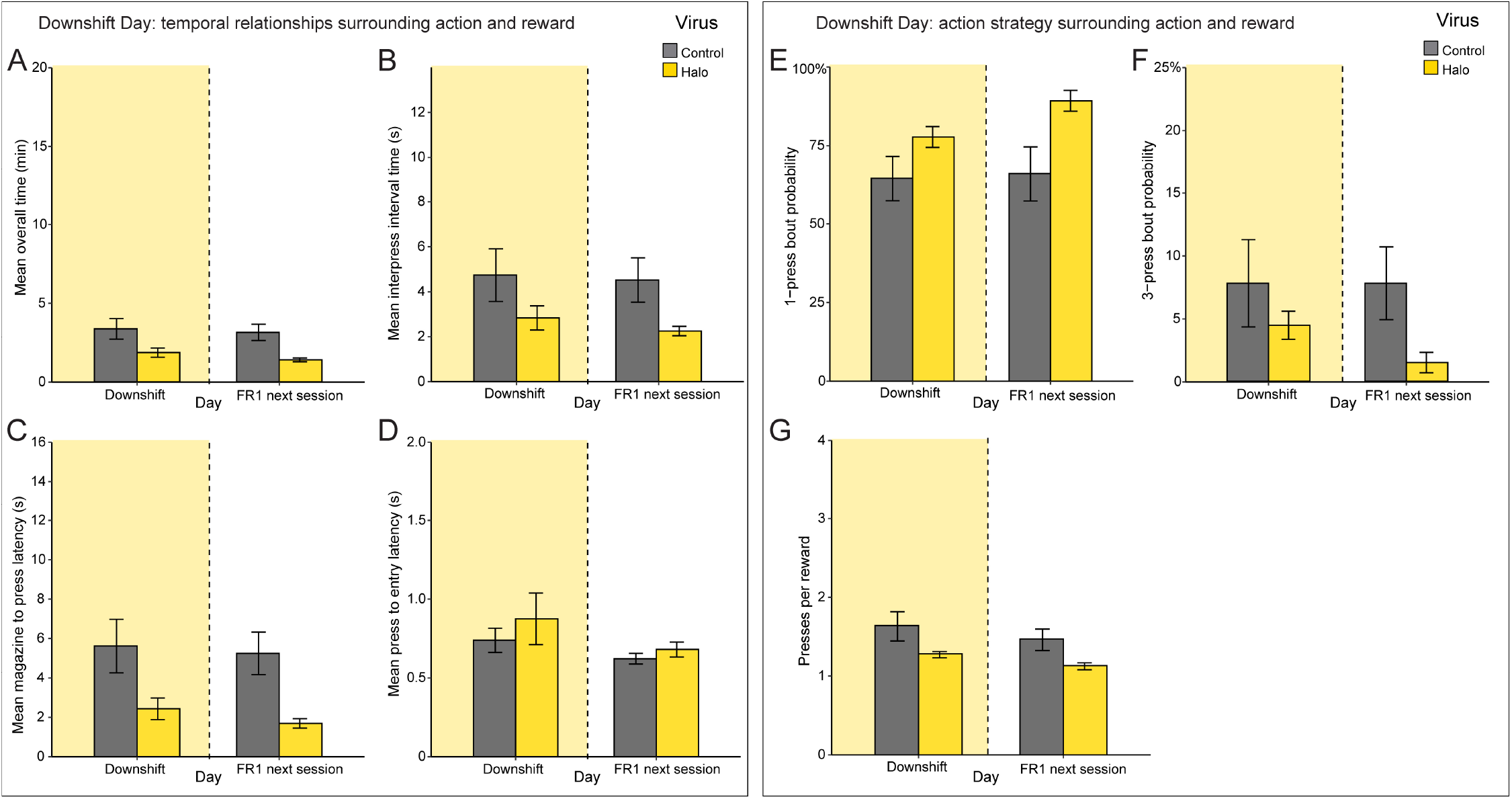
Downshift illumination day. 1-sec continuous DLS illumination on every press (reset by magazine entry). Illumination effects were compared within- and between-groups to a normal FR1 session the day immediately after. **A**, Mean overall time (min). **B**, Mean inter-press intervals (sec). **C**, Mean reward contact (magazine) to press latency (sec). **D**, Mean lever press to a magazine entry (sec). **E**, Mean 1-press bout probability (%). **F**, Mean 3-press bout probability (%). **G**, Total presses per reward (ratio). Control, grey; eNpHR3.0, yellow. For all figure panels, bars and errors show mean ± SEM; asterisks denote significant comparisons (\**P*<0.05, \*\**P*<0.01, \*\*\**P*<0.001); lack of asterisks denotes lack of significance.

#### Action strategy data

The halorhodopsin group performed more 1-press bouts during the illumination day, which continued into the subsequent day as well (Fig. 4E). This was reflected by a main effect of group (estimate: 2.76, CI: 1.16 – 6.56, *p* = 0.022) and a day/group interaction (estimate: 1.51, CI: 1.11 – 2.07, *p* = 0.009), but no day effect (estimate: 1.04, CI: 0.86 – 1.26, *p* = 0.690). There was a trend towards fewer 3-press bouts in halorhodopsin rats (Fig. 4F), but there was not a main effect of day (estimate: 1.00, CI: 0.72 – 1.39, *p* = 1.000), a day/group interaction (estimate: 0.56, CI: 0.29 – 1.09, *p* = 0.091), or a group effect (estimate: 0.37, CI: 0.12 – 1.13, *p* = 0.082).

The total number of presses performed were lower for the halorhodopsin group on the illumination day and subsequent day (Fig. 4G), as seen in a main effect of day (estimate: −0.09, CI: −0.15 – −0.02, *p* = 0.015) and group (estimate: −0.35, CI: −0.64 – −0.05, *p* = 0.021), but no day and group interaction (estimate: 0.01, CI: − 0.09 – 0.11, *p* = 0.822). In this FR1 condition, 1 press per reward would be most optimal, and the halorhodopsin rats were closest to that level.

### RT30 Days

Changes in the timing and frequency of lever-press actions during upshift and downshift days could possibly be related to greater/lesser probability of reward occurring given the action. This is difficult to discern in FR schedules, as pressing directly produces reward delivery. Thus, as a final experimental phase we exposed animals to a shift from FR1 to a RT30 schedule, which is similar to what we had done previously (Amaya et al., 2021). Under RT30, rewards were delivered independently of pressing every ∼30sec. This schedule affords an opportunity to evaluate how changes in lever press behavior might relate to how animals experience the relationships between the action and reward under normal conditions or with DLS inhibition, which we tested for three sequential RT30 days with DLS illumination.

#### Temporal data

Animals with DLS inhibition exhibited a remarkably faster performance rate. Inter-press-intervals expectedly rose in control animals over RT30 days, while these intervals in animals with DLS inhibition remained low (i.e., faster). This was seen in a day by group interaction (estimate: 0.70, CI: 0.52 – 0.0.94, *p* = 0.019) (Fig 5A). Similarly, animals with DLS inhibition showed lower press-to-magazine entry times; day by group interaction; estimate: 0.83, CI: 0.70 – 0.97, *p* = 0.021) (Fig C). They also exhibited a trend towards a shorter time between a magazine entry and continuation of pressing behavior, but the interaction of day and group was not significant (estimate: 0.98, CI: 0.82 – 1.17, *p* = 0.804) (Fig B). In total, the RT30 sessions were characterized in control animals by a slowing of actions, while DLS inhibition caused animals to maintain a relatively speedier performance.

**Figure 5.**
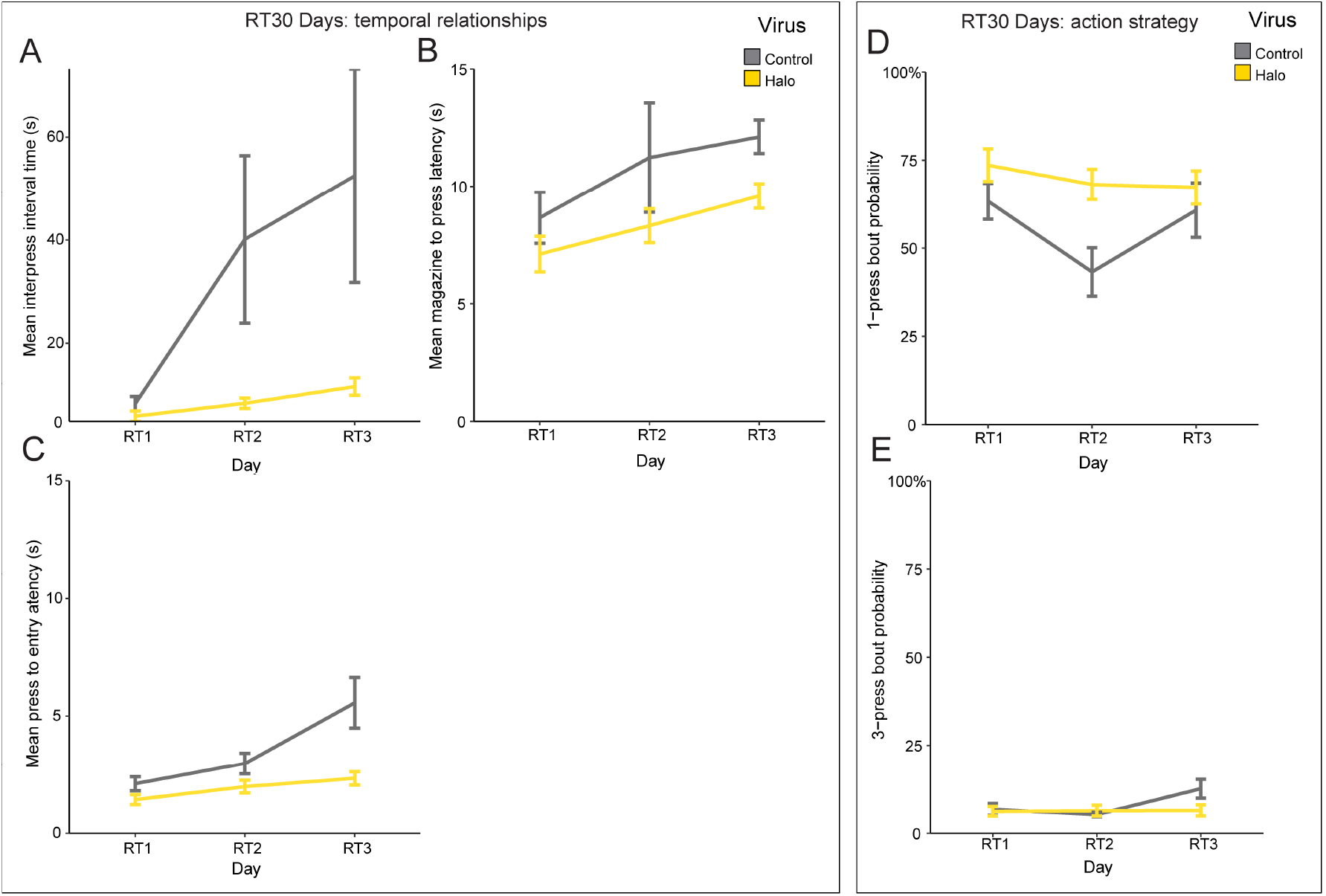
Random time-30 days. 1-sec continuous DLS illumination on every press (no reset by magazine entry). Illumination effects were compared within- and between-groups to a normal FR1 session the day immediately after. **A**, Mean inter-press intervals time (sec). **B**, Mean press-to-magazine entry (sec). **C**, Mean reward contact (magazine) to press latency (sec). **D**, Mean 1-press bout probability (%). **E**, Mean 3-press bout probability (%). Control, grey; eNpHR3.0, yellow. For all figure panels, bars and errors show mean ± SEM; asterisks denote significant comparisons (\**P*<0.05, \*\**P*<0.01, \*\*\**P*<0.001); lack of asterisks denotes lack of significance.

#### Action strategy data

The dominant strategy of behavior in all animals was to continue performing 1-press bouts during RT30 days. Animals with DLS inhibition showed more of these 1-press bout events overall, which did not change across RT30 days (day by group interaction; estimate: 1.22, CI: 1.03 – 1.46, *p* = 0.024) (Fig 5D). The occurrence of 3-press bouts was infrequent, and was not different between experimental groups (estimate: 0.92, CI: 0.67 – 1.25, *p* = 0.577) (Fig 5E). Thus, DLS inhibition led to more of a continuation of past 1-bout pressing strategies, while also speeding up performance.

We also looked at the probability of a lever press action just prior to, and just after, reward delivery. Probabilities were individually set per animal as a rounded mean inter-press interval over the session. In both groups, actions were more likely to occur prior to reward delivery than after reward delivery and pressing in both time windows declined over the course of the sessions, as befitting the RT30 schedule. However, animals with DLS inhibition performed significantly more interactions both before and after reward delivery. In analyses, pre-reward action probability (Fig. 6A) did not reveal a group effect (estimate: 0.73, CI: 0.28 – 1.91, *p* = 0.521) but there were significant effects for both session (estimate: 0.52, CI: 0.39 – 0.68, p < 0.001) and the group/session interaction (estimate: 2.01, CI: 1.15 – 3.52, *p* = 0.015). The probability of an action just after reward (Fig. 6B) followed an identical pattern with no group effect (estimate: 0.63, CI: 0.17 – 2.40, *p* = 0.503) but significant effects for both session (estimate: 0.50, CI: 0.37 – 0.68, *p* < 0.001) and the group/session interaction (estimate: 2.17, CI: 1.19 – 3.95, *p* = 0.012). These trends were reflected in overall press rates across RT30 sessions (Fig. 6C). Analyses revealed both a group (estimate: 4.27, CI: 0.32 – 8.22, *p* = 0.034) and session effect (estimate: −2.74, CI: −3.42 – −2.07, *p* < 0.001), but not their interaction (estimate: −0.17, CI: − 1.53 – 1.19, *p* = 0.806). They were also somewhat reflected within the peri-reward time window (+/- 1 second) (Fig. 6D), in which presses per minute was significant for session (estimate: −4.28, CI: −5.94 – −2.62, *p* <0.001), but was not significant for group (estimate: 6.63, CI: −3.60 – 16.86, *p* = 0.204) or the group/session interaction (estimate: 0.33, CI: −2.99 – 3.64, *p* = 0.846).

**Figure 6.**
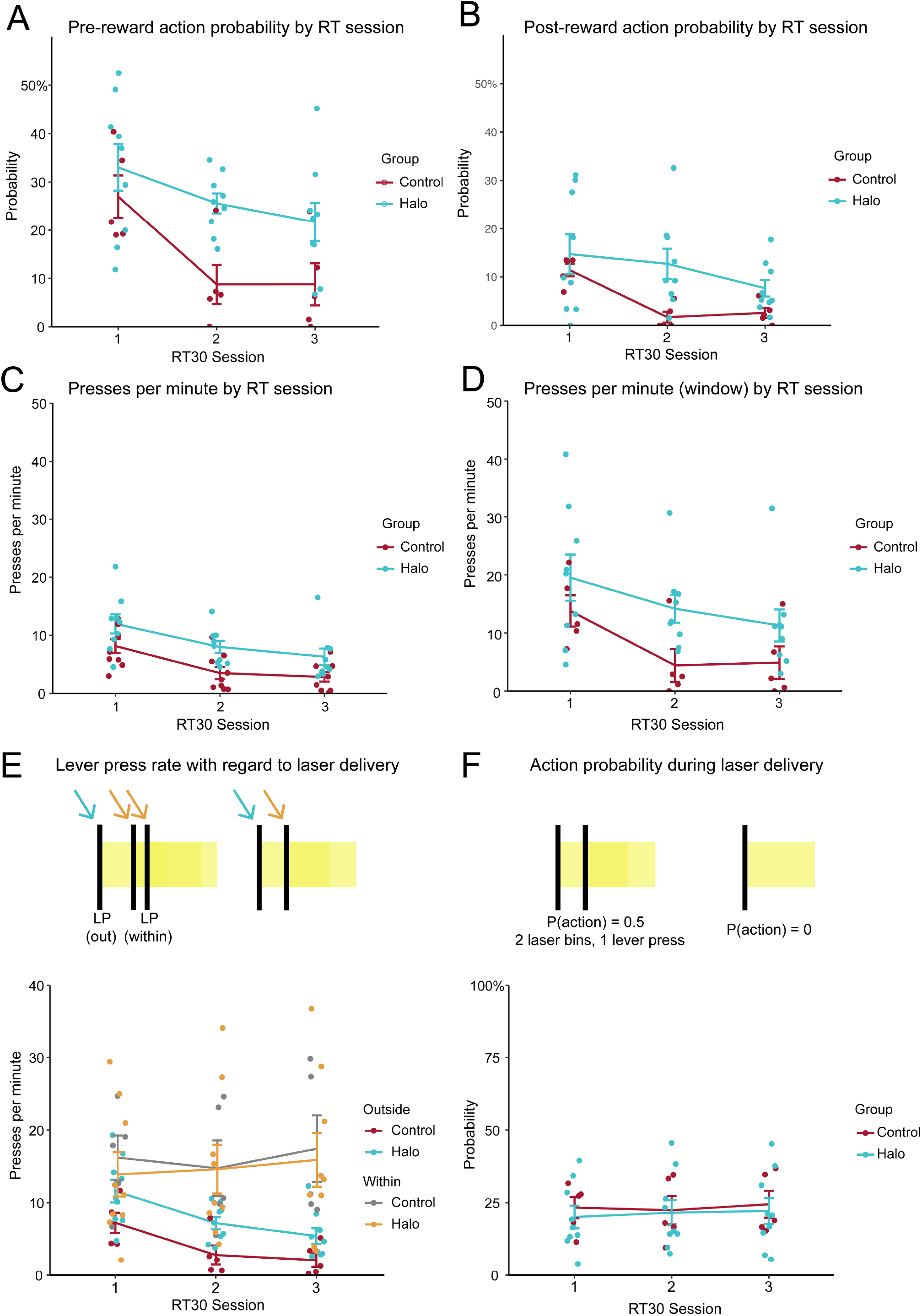
RT30 illumination sessions (3 days). 1-sec continuous DLS illumination on every press (reset by magazine entry). **A**, Pre-reward action probability by RT session. **B**, Post-reward action probability by RT session. **C**, Presses per minute by RT session. **D**, Presses per minute (illumination/laser 1-sec time window) by RT session. **E**, Lever press rate comparison without versus within laser delivery time window. **F**, Action probability during laser delivery; action probability (set per animal as its rounded mean inter-press interval per session).

Further analysis was undertaken for both action probability and lever press rate with regard to DLS illumination timing (Fig. 6E). For this, lever press rates outside the laser delivery time window were considered distinct from those occurring within the light administration time period, and a press that triggered the illumination itself was considered to occur outside of the illumination period. Analyses showed a significant effect of pressing rates between within-illumination time vs. outside-of-illumination time (estimate: −15.47, CI: − 20.87 – −10.08, *p* < 0.001), but no effects were found for group (estimate: 1.28, CI: −4.90 – 7.47, *p* = 0.684), session (estimate: −0.98, CI: −3.07 – 1.11, *p* = 0.357), or any interactions between group and session (estimate: −0.07, CI: −2.67 – 2.54, *p* = 0.960), group and illumination time (estimate: 4.46, CI: −2.27 – 11.19, *p* = 0.194), session and illumination time (estimate: −3.19, CI: −7.37 – 0.99, *p* = 0.134), or a group/session/illumination time interaction (estimate: −0.90, CI: −6.12 – 4.31, *p* = 0.734). Concerning action probability, this was calculated as a lever press occurring during the illumination period (binary: yes/no) as each lever press resulted in the triggering of a laser pulse during the RT session. We did not find any significant effects for group (estimate: 0.78, CI: 0.32 – 1.88, *p* = 0.582), session (estimate: 1.04, CI: 0.85 – 1.27, *p* = 0.690), or a group/session interaction (estimate: 1.03, CI: 0.80 – 1.33, *p* = 0.806) (Fig. 6F). Thus, actions tended to be performed more frequently within the illumination period regardless of experimental group.

These data regarding action probabilities and illumination times indicated no major differences between groups in *when* actions would occur, when they did occur. Although a relatively higher rate and frequency of actions occurred in animals with DLS inhibition, the distribution of those actions was thus no different than controls. This result suggests that differences between groups were unlikely to be related to differences in extra positive reinforcement gained in DLS inhibition animals by virtue of their greater overall action frequency, as these animals distributed their increased pressing before reward as well as after reward (rather than just before reward). Results are similarly unlikely explained by DLS inhibition improving the timing of actions to reflect a ∼30 sec reward delivery schedule; in such a case we would have expected more of a piling up of actions just preceding reward delivery in the DLS inhibition group more so than in controls.

## Discussion

It is advantageous to repeat actions if they have worked well in the past. However, when conditions change and those familiar action plans are no longer optimal, it can be better to learn new ways of behaving. This balance between continuing with old action patterns versus replacing them is in part the domain of the striatum. Curiously, the dorsolateral striatum can promote both new action learning as well as the usage of acquired actions. What does it do when these two options are in conflict when conditions change? We probed this by suddenly changing the task rules on rat subjects and assessing how they react (Amaya & Smith, 2021; Balleine & Killcross, 1994; Balleine et al., 2003; Yin & Knowlton, 2006; Yin et al., 2006). We specifically challenged a well-learned behavior (1-press action bouts during FR1) with changes in action contingencies (e.g., upshift, downshift, RT30) while inhibiting the DLS. During an FR1→FR3 upshift, DLS inhibition caused animals to adapt less readily and stick to the old one-press action strategy more. Performance vigor was reduced as well. Later, during an FR3→FR1 downshift, and then during the FR3-RT30 shift, the same was seen: animals favored using the 1-press bout strategy they initially learned to do. This time, however, performance vigor was increased. We conclude that the DLS inhibition caused animals to use their old action plans. When goal procurement was less compatible with those (upshift), vigor was reduced. When goal procurement was more compatible with those (downshift, RT), vigor was increased. By extension, the results support an interpretation of the DLS as helping promote a prospective form of action learning.

This result diverges from what it is seen with inhibition of specifically the small population of DLS cholinergic interneurons, in which animals are better at adapting their behavior to changes in task rules (Amaya & Smith, 2021). With neuron-wide DLS inhibition here, animals did seemingly the opposite by favoring the initially learned FR1 style of behavior and appearing as more inflexible. Differences in DLS manipulation methods between these studies aside, it could be that cholinergic neurons and non-cholinergic neurons within the DLS carry divergent functions, a possibility that will require additional neuron-specific targeting approaches to resolve.

These results also may diverge from those of a recent study finding that activity in DLS neurons reflected previously experienced maze running experience better than forthcoming running goals (Cunningham et al., 2021), suggesting they represent retrospectively oriented (i.e., previously learned) information. Here, it would appear conversely that prior experience (i.e., FR1-driven behavior) was dominant when the DLS was actually offline. Reconciling these results will require a better understanding of how our indiscriminate DLS manipulation relates to likely cell type-specific functionality as this study shows, and a better understanding of why DLS inhibition was most effective during task change days rather than during behavior without task changes (i.e., FR3 maintenance day).

Factors of potential interest in action flexibility vs. inflexibility here would include behavioral inhibition, positive and negative error learning, incentive motivation, and energy expenditure. The inconsistent effects of DLS inhibition on vigor across the different task shifts makes it unlikely that changes in basic processes such as reward valuation or motor control explain the results. Concerning reward in particular, some DLS neurons respond to reward (Schmitzer-Torbert & Redish, 2004; Smith & Graybiel, 2016). Here, with the dominant overall trend on illumination test days being to perform 1-press bouts, the 1-sec illumination wound up extending slightly into a post-press magazine entry for some animals, and was a common occurrence during the downshift day when time between a press and magazine entry averaged less than 1 sec. On this downshift day, animals would typically be encountering reward during this magazine entry (i.e., it was an FR1 schedule). On upshift or maintenance days, animals would not be encountering reward (i.e., they were FR3 schedules). In one view, this could have affected reward information that somehow resulted in animals favoring the FR-1 style of behavior each time (notwithstanding the inconsistent effect on performance speeds). However, the effect of DLS inhibition on behavior was rather different on the upshift vs. maintenance days, with both occurring during an FR3 schedule. Also, during the RT30 day, we were able to see that animals with DLS inhibition did not have a different distribution of their lever press actions around reward delivery, which might have occurred if reward-related information were being affected by DLS inhibition.

DLS inhibition also had markedly different effects on measures of performance vigor depending on the task conditions. Animals were predictably slower during the FR1-FR3 upshift with an inhibited DLS. Such changes in performance vigor can be evident as animals simply perform a learned behavior, but we have found that vigor changes resulting from DLS inhibition are far more pronounced when there is a change in task requirements, specifically a devaluation of the task outcome (Crego et al., 2016; Crego et al., 2020). We note that similarly robust changes in task performance time were not observed during the FR3 maintenance day, in which there was no task shift but in which the DLS was identically inhibited. It begins to get curious then how it is that animals with DLS inhibition were instead *faster* during the FR3-FR1 downshift and also during the RT30 shift. In other words, why would the same DLS inhibition in the same animals reduced performance time during one task change (upshift) while increasing performance time during other task changes (downshift and RT30)? One answer may lie in the fact that the task demands were increased during the upshift (i.e., more actions required for reward), while they were in principle decreased during the downshift and RT30 days (i.e., fewer actions required for reward). In this way, normally the DLS promotes increased vigor, particularly when action demands increase (upshift), but can also encourage decreased vigor when task demands decrease (downshift; RT30).

With a few exceptions, changes in performance vigor tended to occur when animals were moving to a lever press (i.e., the onset of an action), whether the prior event was another press or a magazine entry. By contrast, less consistent speed changes were seen at the plausible time of action termination (e.g., from a lever press to a magazine entry). This result is consistent with findings showing that DLS activity, and the activity of its dopaminergic input, at the onset of an action sequence carries a great deal of control over how vigorous that action sequence is (Barnes et al., 2005; Crego et al., 2020; da Silva et al., 2018; Jin & Costa, 2010; Jog et al., 1999; Smith & Graybiel, 2013a; Thorn et al., 2010).

We cannot touch on the relation of our data to cell-type-specific functions of the DLS. Electrophysiological studies show a considerable degree of heterogeneity concerning how and when different DLS neurons respond during habitual behaviors (e.g., (Smith & Graybiel, 2013a)). Molecular-targeting approaches have created opportunities to better understanding two distinctions of cell type, those being DLS projection cells in the direct and indirect basal ganglia pathways. Although the two pathways are thought to have opposing roles in movement, there can be synchronization or cooperation happening between them for action initiation and habits (Cui et al., 2013; Isomura et al., 2013; Jin, et al., 2014; O’Hare et al., 2016; Tecuapetla et al., 2016). Separately, evidence is accumulating that the indirect pathway within the DLS is particularly critical for habitual expression of behavior (Corbit et al., 2014; Smith et al., 2021; Vicente et al., 2016; Yu et al., 2009). Here, perhaps DLS inhibition created a roughly synchronous deactivation of neurons in both pathways, which could, depending on task state, create different types of consequences on behavior depending on whether the pathways would normally be cooperating, independent, or competing in function. It is relevant that these pathways show divergent functions in the dorsomedial striatum for ongoing behavior depending on whether task demands are simple or more challenging (Bolkan et al., 2021), raising the possibility that cognitive demand in FR1 vs. FR3 conditions differentially engage these pathways.

Our results point to a larger challenge to the study of actions and habits. Typically, habits are inferred from a negative result (i.e., lack of sensitivity to reward devaluation or action-outcome contingencies). Although measures of vigor have proven to be a helpful positive identifier of habits, one can also find goal-directed behaviors that look just as skillful as habits (Balleine et al., 2009; Vandaele & Janak, 2023; Garr & Dalamater, 2019). Recently, reward devaluation studies have uncovered problems associated with using this method to determine a behavior as a habit, as the context of the devaluation procedure itself carries great weight over observed results (Amaya et al., 2020; Bouton et al., 2021). Looking in detail at *how* animals behave in different task conditions, such as the vigor and bout-type aspects here, may help create new positive identifiers of habits (Amaya & Smith, 2018; Elzelingen et al., 2022; Graybiel, 2008; Lerner, 2020; Malvaez & Wassum, 2018; Schreiner et al., 2019; Watson et al., 2022).

## Acknowledgements

We thank Dr. Neil Winterbauer, Dr. Stephen Chang, Dr. Alyssa DiLeo, Alex Brown, and Matthew Betz for assistance. This work was supported by an NSF research grant to KSS (IOS 1557987), an NIH grant to KSS (R01DA04419), and an NIH grant to KAA (F99NS115270).

## References

Amaya KA, Smith KS (2018). Neurobiology of habit formation. Current Opinion in Behavioral Sc iences, 20:145–152.

Amaya KA, Smith KS (2021). Spatially restricted inhibition of cholinergic interneurons in the dorsolateral striatum encourages behavioral exploration. European Journal of Neuroscience, 53(8):2567–2579.

Amaya KA, Stott JJ, Smith KS (2020). Sign-tracking behavior is sensitive to outcome devaluation in a devaluation context-dependent manner: implications for analyzing habitual behavior. Learning & Memory, 27:136–149.

Bailey KR, Mair RG (2006). The role of striatum in initiation and execution of learned action sequences in rats. The Journal of Neuroscience, 26(3):1016.

Balleine BW, Killcross AS (1994). Effects of ibotenic acid lesions of the nucleus accumbens on instrumental action. Behavioural Brain Research, 65:181–193.

Balleine BW, Killcross AS, Dickinson A (2003). The effect of lesions of the basolateral amygdala on instrumental conditioning. The Journal of Neuroscience, 23:666–675.

Balleine BW, Liljeholm M, Ostlund SB (2009). The integrative function of the basal ganglia in instrumental conditioning. Behav Brain Res, 199(1):43–52.

Barker J, Taylor J (2014). Barker JM, Taylor JR. Habitual alcohol seeking: modeling the transition from casual drinking to addiction. Neurosci Biobehav Rev 47:281–294. Neuroscience & Biobehavioral Reviews, 47.

Barnes TD, Kubota Y, Hu D, Jin DZ, Graybiel AM (2005). Activity of striatal neurons reflects dynamic encoding and recoding of procedural memories. Nature, 437(7062):1158–1161.

Bates D, Mächler M, Bolker B, Walker S (2015). Fitting linear mixed-effects models using lme4. Journal of Statistical Software, 67(1):1–48.

Bergstrom, H. C., Lipkin, A. M., Lieberman, A. G., Pinard, C. R., Gunduz-Cinar, O., Brockway, E. T., … Holmes, A. (2018). Dorsolateral striatum engagement interferes with early discrimination learning. Cell Rep, 23(8):2264–2272.

Bolkan S, Stone I, Pinto L, Ashwood Z, Garcia J, Herman A, … Witten I (2021). Strong and opponent contributions of dorsomedial striatal pathways to behavior depends on cognitive demands and task strategy. Nat Neurosci, 25:345–357.

Bouton ME, Allan SM, Tavakkoli A, Steinfeld MR, Thrailkill EA (2021). Effect of context on the instrumental reinforcer devaluation effect produced by taste-aversion learning. Journal of Experimental Psychology: Animal Learning and Cognition, 47:476–489.

Corbit LH, Nie H, Janak PH (2014). Habitual responding for alcohol depends on both AMPA and D2 receptor signaling in the dorsolateral striatum. Font Behav Neurosci, 8:301.

Corbit LH, Janak PH (2016). Habitual alcohol seeking: neural bases and possible relations to alcohol use disorders. Alcoholism: Clinical and Experimental Research:40.

Crego ACG, Chang SE, Butler WE, Smith KS (2016). Optogenetic research in behavioral neuroscience: insights into the brain basis of reward learning and goal-directed behavior. In: Optogenetics: From Neuronal Function to Mapping & Disease Biology. Cambridge University Press.

Crego ACG, Stocek F, Marchuk AG, Carmichael JE, van der Meer MAA, Smith KS (2020). Complementary control over habits and behavioral vigor by phasic activity in the dorsolateral striatum. Journal of Neuroscience, 40(10):2139–2153.

Cromwell HC, Berridge KC (1996). Implementation of action sequences by a neostriatal site: a lesion mapping study of grooming syntax. The Journal of Neuroscience, 16(10):3444.

Cui G, Jun SB, Jin X, Pham MD, Vogel SS, Lovinger DM, Costa RM (2013). Concurrent activation of striatal direct and indirect pathways during action initiation. Nature, 494(7436):238–242.

Cunningham PJ, Regier PS, Redish AD (2021). Dorsolateral striatal task initiation bursts represent past experiences more than future action plans. The Journal of Neuroscience, 41(38):8051–8064.

da Silva JA, Tecuapetla F, Paixao V, Costa RM (2018). Dopamine neuron activity before action initiation gates and invigorates future movements. Nature, 554(7691):244–248.

Dezfouli A, Balleine BW (2012). Habits, action sequences and reinforcement learning. European Journal of Neuroscience, 35(7):1036–1051.

Elzelingen W, Warnaar P, Matos J, Bastet W, Jonkman R, Smulders D, … Willuhn I (2022). Striatal dopamine signals are region specific and temporally stable across action-sequence habit formation. Current Biology, 32.

Garr E, Delamater AR (2019). Exploring the relationship between actions, habits, and automaticity in an action sequence task. Learning and Memory, 26:128–132.

Gasbarri A, Pompili A, Packard M, Tomaz C (2014). Habit learning and memory in mammals: Behavioral and neural characteristics. Neurobiology of learning and memory, 114:198–208.

Goodman J, Ressler RL, Packard MG (2017). Enhancing and impairing extinction of habit memory through modulation of NMDA receptors in the dorsolateral striatum. Neuroscience, 352:216–225.

Graybiel AM (2008). Habits, rituals, and the evaluative brain. Annu Rev Neurosci, 31:359–387.

Isomura Y, Takekawa T, Harukuni R, Handa T, Aizawa H, Takada M, Fukai T (2013). Reward-modulated motor information in identified striatum neurons. Journal of Neuroscience, 33(25):10209–10220.

Jin X, Costa RM (2010). Start/stop signals emerge in nigrostriatal circuits during sequence learning. Nature, 466(7305):457–462.

Jin X, Tecuapetla F, Costa RM (2014). Basal ganglia subcircuits distinctively encode the parsing and concatenation of action sequences. Nat Neurosci, 17(3):423–430.

Jog MS, Kubota Y, Connolly CI, Hillegaart V, Graybiel AM (1999). Building neural representations of habits. Science, 286(5445):1745–1749.

Knowlton B, Patterson T (2018). Habit formation and the striatum. Curr Top Behav Neurosci. 37:275–295.

Kuznetsova A, Brockhoff PB, Christensen RHB (2017). lmerTest package: tests in linear mixed effects models. Journal of Statistical Software, 82(13):1–26.

Lerner TN (2020). Interfacing behavioral and neural circuit models for habit formation. J Neurosci Res, 98(6):1031–1045.

Lovinger DM, Gremel CM (2021). A circuit-based information approach to substance abuse research. Trends in Neurosciences, 44:122–135.

Malvaez M, Wassum KM (2018). Regulation of habit formation in the dorsal striatum. Current Opinion in Behavioral Sciences, 20:67–74.

O’Hare JK, Ade KK, Sukharnikova T, Van Hooser SD, Palmeri ML, Yin HH, Calakos N (2016). Pathways-specific striatal subtrates for habitual behavior. Neuron, 89(3):472–479.

Packard MG, McGaugh JL (1992). Double dissociation of fornix and caudate nucleus lesions on acquisition of two water maze tasks: Further evidence for multiple memory systems. Behavioral Neuroscience, 106:439–446.

Panigrahi B, Martin KA, Li Y, Graves AR, Vollmer AR, Olson L, Mensh BD, Karpova AY, Dudman Jt (2015). Dopamine Is Required for the Neural Representation and Control of Movement Vigor. Cell, 162(6):1418–1430.

Regier PS, Amemiya S, Redish AD (2015). Hippocampus and subregions of the dorsal striatum respond differently to a behavioral strategy change on a spatial navigation task. J Neurophysiol. 114(3):1399–1416.

Schmitzer-Torbert N, Redish AD (2004). Neuronal activity in the rodent dorsal striatum in sequential navigation: separation of spatial and reward responses on the multiple T task. Journal of Neurophysiology, 91(5):2259–2272.

Schreiner D., Renteria R, Gremel C (2019). Fractionating the all-or-nothing definition of goal-directed and habitual decision-making. J Neurosci Res:98.

Seger CA (2018). Corticostriatal foundations of habits. Current Opinion in Behavioral Sciences, 20, 153–160.

Smith ACW, Jonkman S, Difeliceantonio AG, O’Connor RM, Ghoshal S, Romano MF, … Kenny OJ (2021). Opposing roles for striatonigral and striatopallidal neurons in dorsolateral striatum in consolidating new instrumental actions. Nature Communications, 12(1):5121.

Smith KS, Graybiel AM (2013a). A dual operator view of habitual nehavior reflecting cortical and striatal dynamics. Neuron, 79(3):608–608.

Smith KS, Graybiel AM (2013b). Using optogenetics to study habits. Brain Res, 1511:102–114.

Smith KS, Graybiel AM (2016). Habit formation coincides with shifts in reinforcement representations in the sensorimotor striatum. Journal of Neurophysiology, 115(3):1487–1498.

Tecuapetla F, Jin X, Lima SQ, Costa RM (2016). Complementary contributions of striatal projection pathways to action initiation and execution. Cell, 166(3):703–715.

Thorn CA, Atallah H, Howe M, Graybiel M (2010). Differential dynamics of activity changes in dorsolateral and dorsomedial striatal loops during learning. Neuron, 66(5):781–795.

Tolman EC (1932). Purposive behavior in animals and men. London, England: Century/Random House UK.

Turner KM, Svegborn A, Langguth M, McKenzie C, Robbins TW (2022). Opposing roles of the dorsolateral and dorsomedial striatum in the acquisition of skilled action sequencing in rats. The Journal of Neuroscience, 42(10):2039.

Vandaele Y, Janak PH (2013). Lack of action monitoring as a prerequisite for habitual and chunked behavior: Behavioral and neural correlates. iScience, 26(1):105818.

Vicente AM, Galvao-Ferreira P, Tecuapetla F, Costa RM (2016). Direct and indirect dorsolateral striatum pathways reinforce different action strategies. Current Biology, 26(7):R267–R269.

Watson P, O’Callaghan C, Perkes I, Bradfield L, Turner K (2022). Making habits measurable beyond what they are not: a focus on associative dual-process models. Neuroscience and Biobehaioral Reviews, 142:104869.

Yin HH, Knowlton BJ (2004). Contributions of striatal subregions to place and response learning. Learn Mem, 11(4):459–463.

Yin HH, Knowlton BJ (2006). The role of the basal ganglia in habit formation. Nature Reviews Neuroscience, 7(6):464–476.

Yin HH, Knowlton BJ, Balleine BW (2004). Lesions of dorsolateral striatum preserve outcome expectancy but disrupt habit formation in instrumental learning. European Journal of Neuroscience, 19(1):181–189.

Yin HH, Knowlton BJ, Balleine BW (2006). Inactivation of dorsolateral striatum enhances sensitivity to changes in the action-outcome contingency in instrumental conditioning. Behav Brain Res, 166(2), 189–196.

Yin HH, Mulcare SP, Hilario MR, Clouse E, Holloway T, Davis MI., … Costa RM (2009). Dynamic reorganization of striatal circuits during the acquisition and consolidation of a skill. Nat Neurosci, 12(3):333–341.

Yttri EA, Dudman JT (2016). Opponent and bidirectional control of movement velocity in the basal ganglia. Nature, 533(7603):402–406.

Yu C, Gupta J, Chen JF, Yin HH (2009). Genetic deletion of A2A adenosine receptors in the striatum selectively impairs habit formation. Journal of Neuroscience, 29(48):15100–15103.

